# Genetic interactions among *Ghd7*, *Ghd7.1*, *Ghd8* and *Hd1* contribute to large variation in heading date in rice

**DOI:** 10.1101/485623

**Authors:** Bo Zhang, Haiyang Liu, Feronique Qi, Zhanyi Zhang, Qiuping Li, Zhongmin Han, Yongzhong Xing

## Abstract

Several major heading date genes are sensitive to photoperiod and jointly regulate heading date in rice. However, it is not clear how these genes coordinate rice heading. In this study, a near-isogenic F_2_ population with *Ghd7*, *Ghd7.1*, *Ghd8* and *Hd1* segregating in the Zhenshan 97 (ZS97) background was used to evaluate the genetic interactions among these four genes under natural long-day (NLD) and natural short-day (NSD) conditions, and a series of *Ghd7.1*-segregating populations in different backgrounds were developed to estimate the genetic effects of *Ghd7.1* on heading date under both conditions. Tetragenic, trigenic and digenic interactions among these four genes were observed under both conditions. In the functional *Hd1* backgrounds, the strongest digenic interaction was *Ghd7* by *Ghd8* under NLD conditions but was *Ghd7* by *Ghd7.1* under NSD conditions. Interestingly, *Ghd7.1* acted as a flowering suppressor under NLD conditions, while it functioned alternatively as an activator or a suppressor under NSD conditions depending on the status of the other three genes. Based on the performances of 16 homozygous four-gene combinations, a positive correlation between heading date and yield was found under NSD conditions, but changed to a negative correlation when heading date was over 90 days under NLD conditions. These results demonstrate the importance of genetic interactions in the rice flowering regulatory network and will help breeders to select favorable combinations to maximize rice yield potential for different ecological areas.

## Introduction

Heading date, a crucial trait for rice expansion to high latitudes, is determined by both genetic factors and environmental cues (Andres & Coupland, 2012). Cultivars with an appropriate heading date will be conductive to high grain yield by fully utilizing the light and temperature resources in their growing regions (Zhang *et al.*, 2015a).

In the last two decades, dozens of quantitative trait loci (QTLs) for rice heading date have been cloned by using biparental populations, germplasm resources and mutants with forward- or reverse-genetics approaches (Yamamoto *et al.*, 2012; Hori *et al.*, 2016; Yano *et al.*, 2016). Among these genes, several major QTLs, especially those cloned from natural variations, have pleiotropic effects on heading date, plant height and grain yield, which have been widely subjected to artificial selection in the process of rice genetic improvement. For example, *Heading date1* (*Hd1*), the homolog of *Arabidopsis CONSTANS* (*CO*), encodes a zinc finger CCT (CO, CO-LIKE and TIMING OF CAB1) domain and acts as a major flowering activator in rice (Yano *et al.*, 2000; Zhang *et al.*, 2017). *Hd1* delays heading date in some varieties under long-day (LD) conditions by interacting with other flowering genes such as *Ghd7,* resulting in a taller plant and more grain yield (Nemoto *et al.*, 2016; Zhang *et al.*, 2017). *Ghd7* is a rice-specific gene encoding a CCT domain protein and is important for heading date, grain yield, rice adaptation and drought resistance (Xue *et al.*, 2008; Weng *et al.*, 2014). Another major QTL, *Ghd8* (allelic to *Hd5* and *DTH8*), encodes a HAP3 subunit of heterotrimeric heme activator protein (HAP) and simultaneously controls heading date, plant height and grain number (Wei *et al.*, 2010; Yan *et al.*, 2011; Fujino *et al.*, 2013). *Ghd7.1*, allelic to *DTH7*, *OsPRR37* and *Hd2* and encoding a PSEUDORESPONSE REGULATOR 7-like protein harboring the CCT domain, greatly represses heading and increases grain yield under LD conditions (Koo *et al.*, 2013; Yan *et al.*, 2013; Gao *et al.*, 2014). Natural variations in *OsPRR37/Ghd7.1* also contribute to rice cultivation at a wide range of latitudes (Koo *et al.*, 2013; Yan *et al.*, 2013). It was initially demonstrated that these genes are in separate branches in the flowering regulatory network and have partially unrelated effects on transcription level (Brambilla & Fornara, 2013; Song *et al.*, 2015).

Photoperiod sensitivity largely determines heading date in rice. There are two independent genetic pathways involved in photoperiod sensitivity. One is the *OsGI-Hd1-Hd3a* pathway, which is conserved with the *GI-CO-FT* pathway in *Arabidopsis* (Shrestha et al., 2014). *Hd1* is upregulated by *OsGI* and activates the expression of *Hd3a* to promote rice heading under both short-day (SD) and LD conditions (Hayama *et al.*, 2003; Zhang *et al.*, 2017). Another is the *Ehd1-Hd3a* pathway, a unique pathway in rice regulated by many genes (Doi *et al.*, 2004; Tsuji *et al.*, 2011). Among these genes, *Ehd2*, *Ehd3*, *Ehd4* and *OsMADS51* always promote rice heading by directly or indirectly upregulating the expression of *Ehd1* under both SD and LD conditions (Kim *et al.*, 2007; Matsubara *et al.*, 2008; Matsubara *et al.*, 2011; Gao *et al.*, 2013). In contrast, other genes including *Ghd7*, *Ghd8*, *Ghd7.1*, *Hd16*, *OsCOL4* and *OsCOL10* repress the expression of *Ehd1*, resulting in late flowering under LD conditions (Xue *et al.*, 2008; Lee *et al.*, 2010; Yan *et al.*, 2011; Hori *et al.*, 2013; Yan *et al.*, 2013; Tan *et al.*, 2016). The recent finding that the Ghd7-Hd1 complex represses *Ehd1* by binding to a cis-regulatory region in the *Ehd1* 5’-UTR suggested that *Hd1* was integrated into the rice-specific genetic pathway (Nemoto *et al.*, 2016).

Our previous studies indicated that *Ghd7* and *Ghd8* in the ZS97 background greatly delayed heading date (non-heading) under NLD conditions because of the presence of *Hd1*, indicating a strong genetic interaction among *Ghd7, Ghd8* and *Hd1* (Zhang *et al.*, 2015a). Ghd7.1 shared the conserved CCT domain with Hd1 and Ghd7, and formed a heterotrimer with Ghd8 and NF-YCs similar to Hd1 (Zhang *et al.*, 2015b; Goretti *et al.*, 2017). Thus, we hypothesized that *Ghd7.1* is involved in genetic interactions with the three other genes. To test this hypothesis, we further conducted genetic interaction analysis among *Ghd7*, *Ghd7.1*, *Ghd8* and *Hd1* in the ZS97 background under NLD and NSD conditions in this study. Tetragenic, trigenic and digenic interactions among these four genes were observed under both NLD and NSD conditions. *Ghd7.1* always acts as a flowering suppressor under NLD conditions but exhibits an alternative function (either suppression or activation) in heading date under NSD conditions.

## Materials and methods

### Construction of NILs and segregating populations

ZS97 carries a functional allele of *Hd1* but a nonfunctional allele of *ghd7*, *ghd7.1* and *ghd8* (Xue *et al.*, 2008; Yan *et al.*, 2011; Yan *et al.*, 2013; Zhang *et al.*, 2017). We previously developed a near-isogenic line (NIL-1) pyramiding functional *Ghd7*^*MH63*^ and *Ghd8*^*9311*^ in a ZS97 background (Zhang *et al.*, 2015a). Another near-isogenic line (NIL-2) in the ZS97 background, which harbored functional *Ghd7.1*^*TQ*^ and nonfunctional *hd1*^*TQ*^ derived from Teqing (TQ), was crossed with NIL-1. Therefore, NIL-F_1_ plants carried heterozygous *Ghd7*, *Ghd7.1*, *Ghd8* and *Hd1* (Fig. S1a; Table S1). The NIL-F_2_ population was developed by self crossing a NIL-F_1_ plant that was genotyped by using the RICE6K SNP array (Fig. S1b) (Yu *et al.*, 2014). To avoid genetic background noise, a NIL-F_2_ individual harboring heterozygous alleles at all four of these genes was used to produce a NIL-F_3_ population with segregation of these four genes by self-pollination. All individuals of the NIL-F_2_ and NIL-F_3_ populations were genotyped at these four gene loci. According to the genotypes of the NIL-F_3_ population, 8 NIL-F_3_ plants, each carrying heterozygous *Ghd7.1* but with different homozygous combinations of the other three genes, were used to generate 8 NIL-F_4_ populations for estimating the genetic effects of *Ghd7.1*. Sixteen NIL-F_3_ plants with different homozygous four-gene combinations were selected to generate 16 four-gene homozygous lines for evaluating yield performance.

### Field experiments and growth conditions

Rice seeds were sown in a seedling bed in the middle of May at the experimental station of Huazhong Agricultural University, Wuhan, China (30.5°N). The 25-day-old seedlings were transplanted into the field with a distance of 16.5 cm between plants within a row and 26.5 cm between rows. The plants were subsequently grown in the field under NLD conditions (a day length of more than 13.5 hours) until the beginning of August. For the field experiments under NSD conditions, the plant materials were sown in Lingshui, Hainan (18.5°N), at the beginning of December and were transplanted into the field after 1 month, at the same planting density as that used in Wuhan, and grown under an average day length of less than 12.5 hours from December to April. The monthly average day length of the growing season in Wuhan and Lingshui is available in Table S2.

The NIL-F_2_ population consisting of 680 individuals was grown in Wuhan in 2016. Excluding the marginal plants and abnormally growing individuals, 509 individuals were used for analysis of genetic interactions among *Ghd7*, *Ghd7.1*, *Ghd8* and *Hd1* under NLD conditions. A total of 900 NIL-F_3_ plants derived from an F_2_ individual segregating for these four genes were grown in Lingshui from Dec 2016 to Apr 2017, and a total of 679 nonmarginal individuals were used for analysis of genetic interactions among these four genes under NSD conditions. Eight NIL-F_4_ populations were grown in Wuhan (~60 plants per population) in summer 2017 (from May to October) and in Lingshui (~40 plants per population) in winter (from Dec 2017 to Apr 2018). Meanwhile, 16 four-gene homozygous lines were also grown in Wuhan and Lingshui in summer and winter of 2017, respectively. Three additional *Ghd7.1*-segregating population (~80 plants per population) with the backgrounds *Ghd7Ghd8Hd1*, *Ghd7Ghd8hd1* and *Ghd7ghd8Hd1* were also grown in Lingshui in winter 2017. In addition, four plants of each four-gene homozygous combination were grown in the field to implement a short-day treatment with a day length of 11 hours and darkness of 13 hours in the summer of 2018. A set of plants from these genotypes were planted in the same field at the same density under NLD conditions and served as the control group.

### DNA extraction, polymerase chain reaction and genotyping

At the tillering stage, leaf blades were collected for DNA extraction using a modified cetyl-trimethyl ammonium bromide (CTAB) method (Murray & Thompson, 1980). Genomic DNA was amplified using rTaq polymerase from TaKaRa in Buffer I according to the manufacturer's indications. For each PCR reaction, DNA was initially incubated for five minutes at 95°C, followed by 35 cycles of amplification (95°C for 30 seconds, 58°C for 30 seconds and 72°C for 30 seconds). The simple sequence repeat (SSR) marker MRG4436, which is tightly linked to *Ghd7*, and the functional markers InDel37, Z9M and S56 designed from *Ghd7.1*, *Ghd8* and *Hd1* (Table S1), respectively, were used to genotype the individuals of all populations and NILs. All markers used for genotyping are listed in Table S5.

### RNA extraction and qRT-PCR analysis

Seedlings were grown in a seedbed under NLD conditions for 30 days and were subsequently transplanted to a plot in the field for the short-day treatment (started on the 11^th^ of June, light treatment from 7:00 am to 6:00 pm every day). After treatment for 15 days (from the 11^th^ to 26^th^ of June), the distal part of young leaves in the short-day treatment and control group (treated with LD condition, i.e., more than 14 hours day length per day from the 11^th^ to 26^th^ of June) were collected at 9:00 am for RNA extraction. For each genotype, leaves from three different individuals were collected as biological replicates. Total RNA was extracted using TRIzol reagent (TransGen Biotech, Beijing) and treated with DNase I (Invitrogen, USA). cDNA was synthesized from 3 μg of RNA using SuperScript III Reverse Transcriptase (Invitrogen, USA). The quantitative analysis of gene expression was performed with SYBR Premix ExTaq reagent (TaKaRa, Dalian) on the ABI ViiA7 Real-time PCR System (Applied Biosystems, USA). The data were analyzed using the relative quantification method. The primers used for real-time PCR are listed in Table S5.

### Trait measurement and data analysis

Heading date was individually scored as the number of days from sowing to the emergence of the first panicle on the plant. The total number of spikelets per plant was measured by the Yield Traits Scorer (Yang *et al.*, 2014). The number of spikelets per panicle (SPP) of each homozygous combination line was recorded as the total number of spikelets divided by the number of panicles. The comparison between genotypes was performed by Student’s *t*-test. To verify the existence of high order genetic interactions, a three-way ANOVA was performed under the condition of fixation of the allele at the fourth gene. The statistical significance of three-way interactions was evaluated by a general liner model (GLM) using the program STATISTICA 8.0 (Statsoft, 1995).

## Results

### The genetic interactions among *Ghd7*, *Ghd7.1* and *Ghd8* under NLD conditions

The NIL-F_1_ plant carrying heterozygous alleles at these four genes (Fig. S1a) was genotyped by the RICE6K SNP array. More than 90% of the NIL-F_1_ plant background was consistent with ZS97 (Fig. S1b). In the NIL-F_2_ population of 509 plants, large variation in heading date was observed, ranging from 65 days to no heading after 160 days under NLD conditions (Fig. S2a). For convenience, 160 days was recorded as the heading date of these non-heading plants. Two-way ANOVA showed that all 6 pairs of digenic interactions of these four genes were highly significant (Table S3). Three-way ANOVA showed that all 4 trigenic interactions were highly significant (Table S4). Four-way ANOVA revealed that the tetragenic interaction among these four genes was also highly significant (*P*<1.0E-10). To better understand the four-way interaction, we classified the populations into three subpopulations based on *Hd1* genotypes: homozygous *Hd1*, heterozygous *Hd1* (*Hd1*^*H*^) and homozygous *hd1*. A significant three-way interaction was detected among *Ghd7*, *Ghd7.1* and *Ghd8* at *P*>1.0E-10 in both the *Hd1* and *Hd1*^*H*^ backgrounds and at *P*=6.9E-04 in the *hd1* background (Fig. S2b-d; Table 1). Additionally, all digenic interactions were detected among *Ghd7*, *Ghd7.1* and *Ghd8.* The *Ghd7* by *Ghd8* interaction contributed more to heading date variation than the other digenic interactions. The square of this interaction accounted for 5.9% and 5.8% of the total sum-of-squares in the *Hd1* and *Hd1*^*H*^ backgrounds, respectively, and 5.8% of that in the nonfunctional *hd1* background (Table 1). The main effects of *Ghd7*, *Ghd8* and their digenic interaction effects explained more than 70% of the variation in heading date in both the *Hd1* and *Hd1*^*H*^ backgrounds. The genetic square of *Ghd7.1* accounted for 17.0% of the total sum-of-squares in the *hd1* background, which was much larger than that observed in the *Hd1* and *Hd1^H^* backgrounds (Table 1). Taken together, these results revealed that a strong trigenic interaction existed among *Ghd7*, *Ghd7.1* and *Ghd8* regardless of the genotype of *Hd1*, and the interaction between *Ghd7* and *Ghd8* showed the strongest digenic interaction among these three genes under NLD conditions.

**Table 1.** Three-way ANOVA analysis of *Ghd7*, *Ghd7.1*and *Ghd8* in NIL-F_2_ population by fixing the genotype of *Hd1* under NLD conditions.

| Effect | DF | *hdl*(n=138; <sup>a</sup>74d-128d) |  |  | *Hdl<sup>H</sup>* (n=242; 67d- <sup>b</sup>160d) |  |  | *Hdl*(n=129; 64d-160d) |  |  |
| --- | --- | --- | --- | --- | --- | --- | --- | --- | --- | --- |
|  |  | F | P | G:T (%) | F | P | G:T (%) | F | P | G:T (%) |
| *Ghd7* | 2 | 787.7 | <1.0E-10 | 28.6 | 12145.5 | <1.0E-10 | 29.6 | 9627.4 | <1.0E-10 | 31.9 |
| *Ghd7.1* | 2 | 468.3 | <1.0E-10 | 17.0 | 747.6 | <1.0E-10 | 1.8 | 799.1 | <1.0E-10 | 2.6 |
| *Ghd8* | 2 | 800.5 | <1.0E-10 | 29.1 | 14400.5 | <1.0E-10 | 35.1 | 11278.0 | <1.0E-10 | 37.4 |
| *Ghd7* by *Ghd7.1* | 4 | 9.6 | 1.1E-06 | 0.7 | 116.5 | <1.0E-10 | 0.6 | 239.9 | <1.0E-10 | 1.6 |
| *Ghd7* by *Ghd8* | 4 | 80.1 | <1.0E-10 | 5.8 | 1183.4 | <1.0E-10 | 5.8 | 893.3 | <1.0E-10 | 5.9 |
| *Ghd7.1* by *Ghd8* | 4 | 21.1 | <1.0E-10 | 1.5 | 34.8 | <1.0E-10 | 0.2 | 31.1 | <1.0E-10 | 0.2 |
| *Ghd7* by *Ghd7.1* by *Ghd8* | 8 | 3.7 | 6.9E-04 | 0.5 | 88.5 | <1.0E-10 | 0.9 | 63.8 | <1.0E-10 | 0.8 |

<sup>a</sup> Range of heading date variation; <sup>b</sup> No heading but recorded as 160 days; *Hdl<sup>H</sup>*, heterozygous allele of *Hdl*; DF, degree of freedom; G:T, ratio of the genetic to the total of sum-of-squares.

### The genetic interactions among *Ghd7*, *Ghd8* and *Ghd7.1* under NSD conditions

The variation of heading date in the NIL-F_3_ population (679 plants) exhibited a continuous distribution ranging from 82 days to 124 days (Fig. S3a). Accordingly, all digenic and trigenic interactions (except the *Ghd7.1* by *Ghd8* by *Hd1* interaction) among these four genes were highly significant under NSD conditions (Tables S3, S4). A significant tetragenic interaction was also observed among these four genes (*P*=2.5E-03). Following the analysis performed for NLD conditions, these 679 plants were also classified into 3 classes according to *Hd1* genotypes. Significant interactions were identified among *Ghd7*, *Ghd7.1* and *Ghd8* in the *hd1*, *Hd1*^*H*^ and *Hd1* backgrounds (Fig. S3b-d; Table 2). However, the digenic interactions among these three genes were different from those detected under NLD conditions. The *Ghd7* by *Ghd7.1* interaction contributed much more to heading date variation than the other two digenic interactions in the functional *Hd1* backgrounds, in which the genetic square accounted for 20.3% and 20.4% of the total sum-of-squares in the *Hd1* and *Hd1*^*H*^ backgrounds, respectively (Table 2). Notably, the effect of *Ghd7* on heading date was the strongest one under NSD conditions, explaining 58%, 21.7% and 29.1% of the variation in the *hd1*, *Hd1*^*H*^ and *Hd1* backgrounds, respectively (Table 2). These results indicated that *Ghd7*, *Ghd7.1* and *Ghd8* interacted under NSD conditions and the *Ghd7* by *Ghd7.1* interaction showed the strongest epistatic effect among the digenic interactions in the functional *Hd1* backgrounds.

**Table 2.**
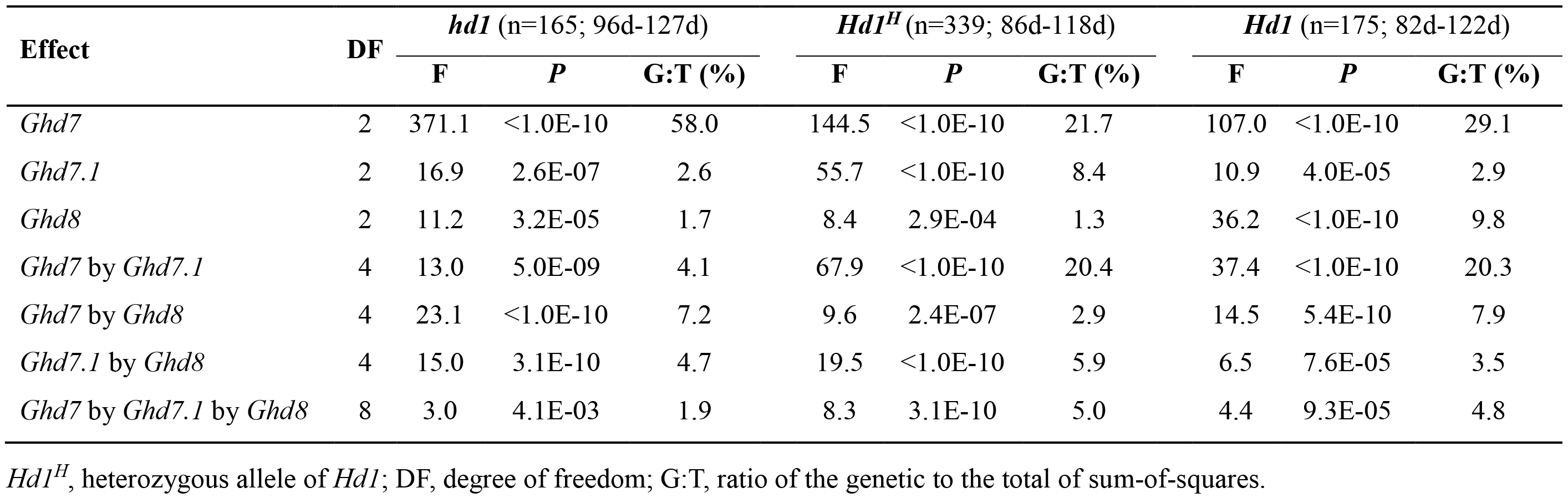
Three-way ANOVA analysis of *Ghd7*, *Ghd7.1*and *Ghd8* in NIL-F_3_ population by fixing the genotype of *Hd1* under NLD conditions.

### *Ghd7.1* as a heading date suppressor under NLD conditions

To estimate the additive and dominance effects of *Ghd7.1* in different genetic backgrounds under NLD conditions, we developed 8 *Ghd7.1*-segregating population (NIL-F_4_) with different homozygous combinations of the other three genes. The NIL-F_4_ population with the *Ghd7Ghd8Hd1* background did not head even after October 24^th^ (the last time when heading date was scored) in Wuhan, when the low temperature is unfavorable to rice growing (Fig. 1a). Therefore, no data were used to evaluate the genetic effect of *Ghd7.1* in this population (Table 3). To confirm whether *Ghd7.1* also delayed rice heading in the *Ghd7Ghd8Hd1* background, we took the young panicles of the main stems of the two homozygous combinations, namely, *Ghd7Ghd7.1Ghd8Hd1* and *Ghd7ghd7.1Ghd8Hd1*, on September 30^th^ and compared their lengths (Fig. 1b, c). The young panicle length of *Ghd7Ghd7.1Ghd8Hd1* (0.87 cm) was significantly shorter than that of *Ghd7ghd7.1Ghd8Hd1* (1.55 cm), which suggested that *Ghd7.1* suppressed heading in the *Ghd7Ghd8Hd1* background under NLD conditions (Fig. 1c). The additive effect of *Ghd7.1* in the other 7 populations ranged from 5.6-19.4 days, indicating that *Ghd7.1* always plays as a suppressor of heading date in these backgrounds (Table 3). The dominance effects and degrees of dominance of *Ghd7.1* ranged from 2.4-10.7 days and from 0.28-0.93, respectively (Table 3).

**Figure 1.**
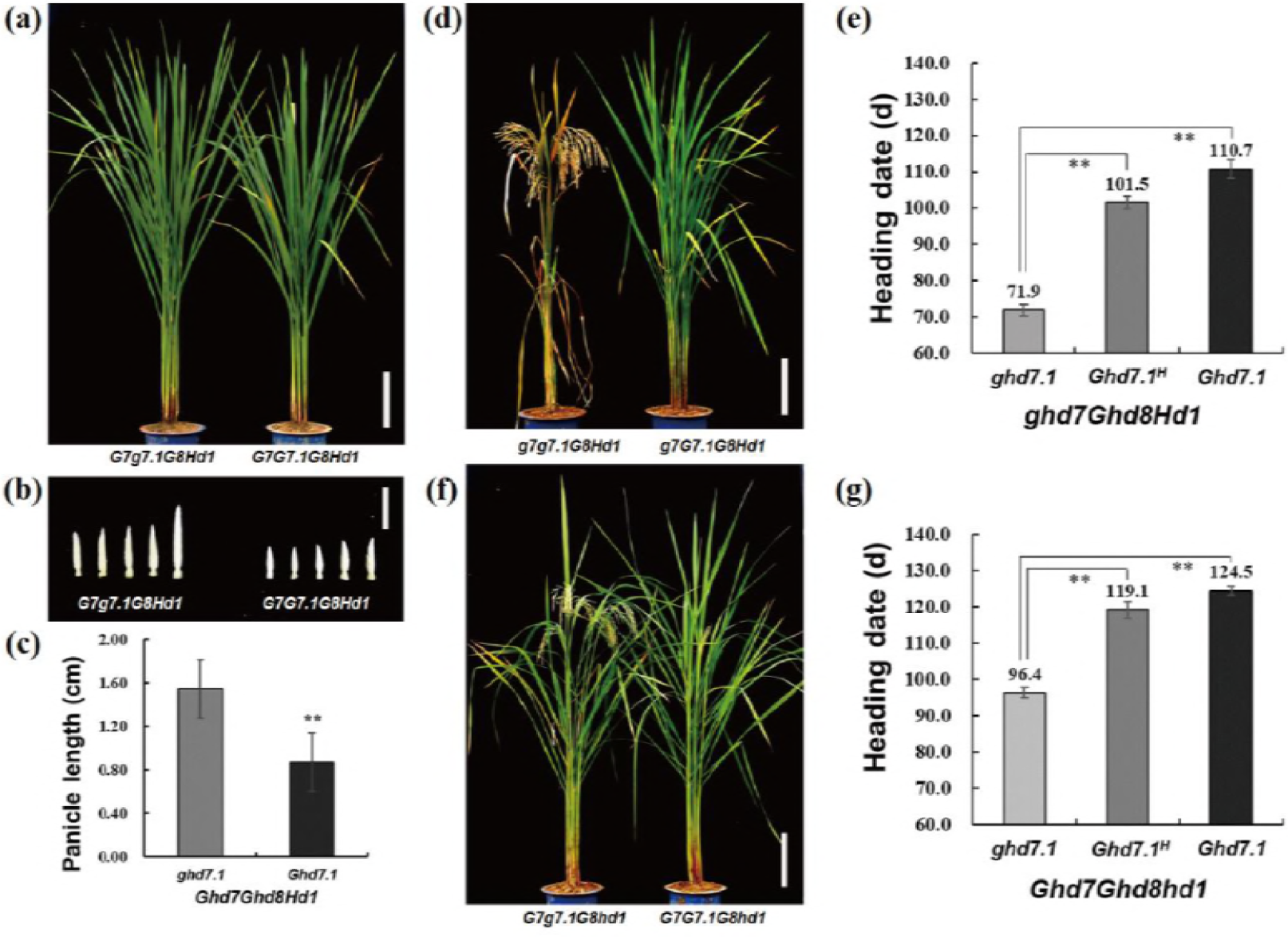
Phenotypes of *Ghd7.1* in different backgrounds under NLD conditions. Non-heading plants of *Ghd7.1* and *ghd7.1* in the functional combination background of *Ghd7*, *Ghd8* and *Hd1* (**a**), their young panicles on main stems taken 136 days after sowing (**b**) and the comparison of panicle lengths (**c**). The plants of *g7G7.1G8Hd1* and *g7g7.1G8Hd1* (**d**); the picture was taken when *g7.1G7G8Hd1* was flowering. The large effect of *Ghd7.1* on heading date in the *ghd7Ghd8Hd1* background (**e**). The plants of *G7G7.1G8hd1* and *G7g7.1G8hd1* (**f**); the picture was taken when *G7G7.1G8Hd1* was flowering. The strong effect of *Ghd7.1* on heading date in the ghd7Ghd8Hd1 background (**g**). **, *P*<0.01 based on Student’s *t*-test; n=15 for each combination in (**c**), and n≥10 for each genotype in (**e**) and (**g**). Scale bars: 20 cm for (**a**), (**d**) and (**f**), and 1 cm for (**b**). *Ghd7.1*^*H*^, the heterozygous allele of *Ghd7.1*.

**Table 3.**
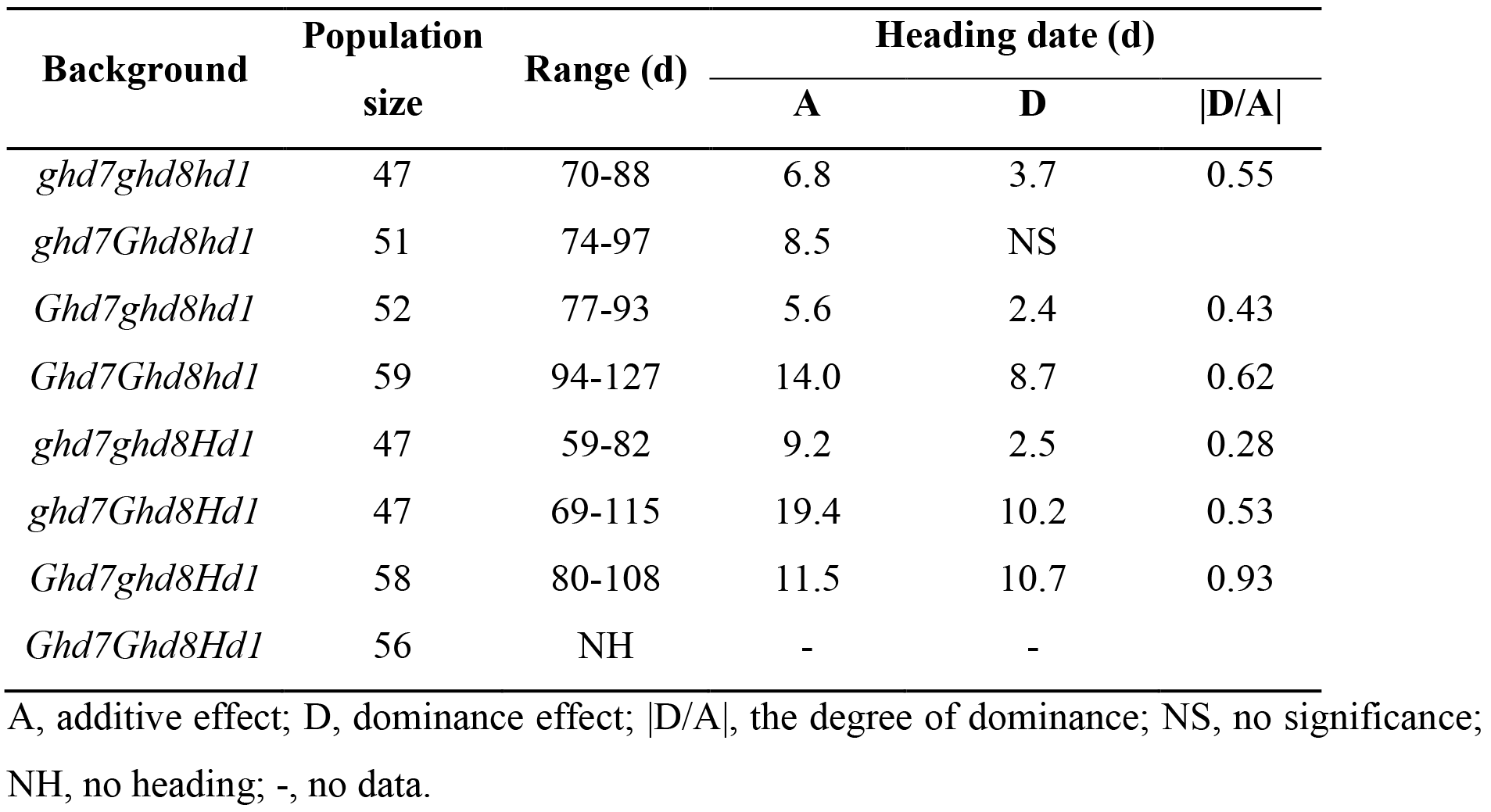
The genetic effects of *Ghd7.1* on heading date in 8 segregating populations with different homozygous combination backgrounds of *Ghd7*, *Ghd8* and *Hd1* under NLD conditions.

| Background | Population size | Range (d) | Heading date (d) |  |  |
| --- | --- | --- | --- | --- | --- |
|  |  |  | A | D | \|D/A\| |
| *ghd7ghd8hd1* | 47 | 70-88 | 6.8 | 3.7 | 0.55 |
| *ghd7Ghd8hd1* | 51 | 74-97 | 8.5 | NS |  |
| *Ghd7ghd8hd1* | 52 | 77-93 | 5.6 | 2.4 | 0.43 |
| *Ghd7Ghd8hd1* | 59 | 94-127 | 14.0 | 8.7 | 0.62 |
| *ghd7ghd8Hd1* | 47 | 59-82 | 9.2 | 2.5 | 0.28 |
| *ghd7Ghd8Hd1* | 47 | 69-115 | 19.4 | 10.2 | 0.53 |
| *Ghd7ghd8Hd1* | 58 | 80-108 | 11.5 | 10.7 | 0.93 |
| *Ghd7Ghd8Hd1* | 56 | NH | - | - |  |

A, additive effect; D, dominance effect; |D/A|, the degree of dominance; NS, no significance; NH, no heading; -, no data.

Additionally, we observed large heading date variations of *Ghd7.1* in the 7 NIL-F_4_ populations, especially in the *ghd7Ghd8Hd1* and *Ghd7Ghd8hd1* backgrounds, in which the variation of heading date ranged from 69-115 days and from 94-127 days, respectively (Table 3). *Ghd7.1* suppressed heading in all these backgrounds. In the *ghd7Ghd8Hd1* background, the difference in heading date between homozygous alleles of *Ghd7.1* and *ghd7.1* was 38.8 days, which was much larger than that in the *ghd7ghd8Hd1* and *ghd7Ghd8hd1* backgrounds (Fig. 1d, e; Table 3), indicating the existence of strong interactions among *Ghd7.1*, *Ghd8* and *Hd1.* In the *Ghd7Ghd8hd1* background, the difference in heading between *Ghd7.1* and *ghd7.1* was 28.1 days, which was also greater than that in the *Ghd7ghd8hd1* and *ghd7Ghd8hd1* backgrounds (Fig. 1f, g; Table 3). These results revealed that the genetic effects of *Ghd7.1* on heading date were dependent on the combinations of *Ghd7*, *Ghd8* and *Hd1.*

### Alternative functions of *Ghd7.1* in repressing or promoting heading under NSD conditions

Under NSD conditions, the additive effects of *Ghd7.1* were 1.8 days, 5.0 days and 2.2 days in the *ghd7ghd8hd1*, *ghd7ghd8Hd1* and *ghd7Ghd8Hd1* backgrounds, respectively (Table 4), indicating that *Ghd7.1* acted as a flowering suppressor in these three backgrounds. However, the effect on delaying heading date was much smaller than that observed under NLD conditions. The genetic effect of *Ghd7.1* disappeared in the *ghd7Ghd8hd1* and *Ghd7ghd8hd1* backgrounds. Interestingly, a converse effect of *Ghd7.1* on heading date was observed in the *Ghd7ghd8Hd1*, *Ghd7Ghd8Hd1* and *Ghd7Ghd8hd1* backgrounds comparing with that observed under NLD conditions. The additive effects of *Ghd7.1* in these three backgrounds were −2.0 days, −9.9 days and −3.7 days, respectively (Table 4). Therefore, it seemed that *Ghd7.1* acted as a heading activator in these three backgrounds. Three additional *Ghd7.1*-segregating populations with the *Ghd7ghd8Hd1, Ghd7Ghd8Hd1* and *Ghd7Ghd8hd1* backgrounds were used to verify this finding. Compared to *ghd7.1, Ghd7.1* promoted rice heading by 4.8 days, 18.0 days and 5.3 days in these three backgrounds, respectively (Fig. 2a-c). In addition, *Ghd7.1* promoted heading by 3.0 days, 12.0 days and 18.2 days in these three backgrounds in the 11-h light treatment, respectively (Fig. 2d-f). These data clearly demonstrsated that *Ghd7.1* acted as a heading activator in the *Ghd7ghd8Hd1*, *Ghd7Ghd8Hd1* and *Ghd7Ghd8hd1* backgrounds under NSD conditions. The dominance effects and degrees of dominance of *Ghd7.1* in these 8 populations ranged from −7.3 to 1.9 days and from 0.38 to 0.88, respectively (Table 4), suggesting that the genetic effects of *Ghd7.1* were largely influenced by the background under NSD conditions.

**Figure 2.**
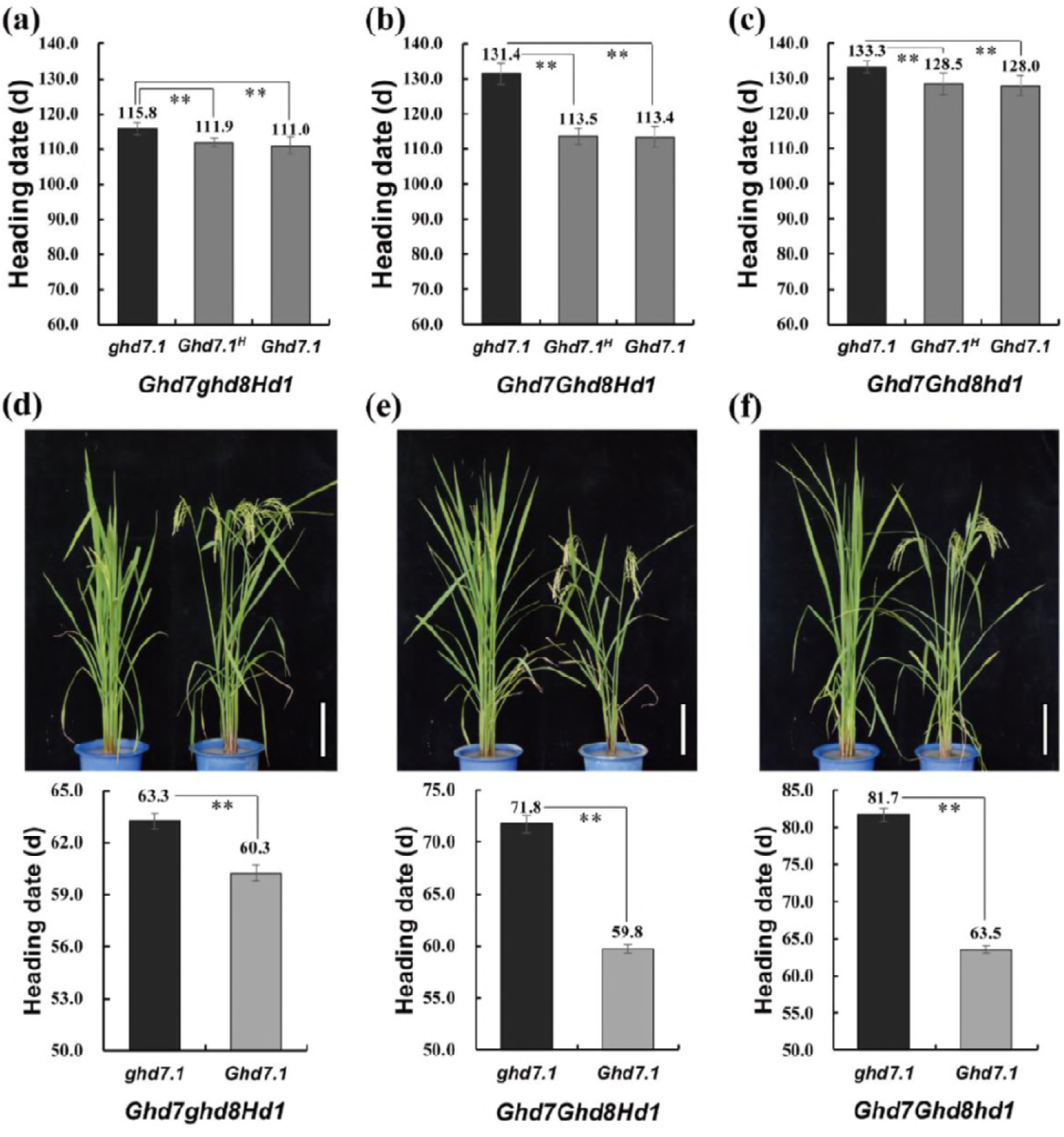
*Ghd7.1* promotes rice heading under NSD and SD conditions. Comparisons of heading date among two homozygotes and a heterozygote of *Ghd7.1* in the backgrounds *Ghd7ghd8Hd1* (**a**), *Ghd7Ghd8Hd1* (**b**) and *Ghd7Ghd8hd1* (**c**) under NSD conditions (n≥10 for each combination). (**d**)-(**f**), Pictures and heading dates of *ghd7.1* and *Ghd7.1* in each corresponding background under SD conditions (11-h light treatment, n=4 for each combination). **, *P*<0.01 based on Student’s *t*-test. *Ghd7.1*^*H*^, the heterozygous allele of *Ghd7.1*. Scale bars: 20 cm for (**d**), (**e**) and (**f**).

**Table 4.** The genetic effects of *Ghd7.1* on heading date in 8 segregating populations with different homozygous combination backgrounds of *Ghd7*, *Ghd8* and *Hd1* under NSD conditions.

| Background | Population size | Range (d) | Heading date (d) |  |  |
| --- | --- | --- | --- | --- | --- |
|  |  |  | A | D | \|D/A\| |
| *ghd7ghd8hd1* | 40 | 112-122 | 1.8 | 1.6 | 0.88 |
| *ghd7Ghd8hd1* | 39 | 110-117 | NS | NS |  |
| *Ghd7ghd8hd1* | 38 | 120-126 | NS | NS |  |
| *Ghd7Ghd8hd1* | 40 | 123-135 | -3.7 | -2.6 | 0.71 |
| *ghd7ghd8Hd1* | 39 | 97-112 | 5.0 | 1.9 | 0.38 |
| *ghd7Ghd8Hd1* | 40 | 102-112 | 2.2 | 1.4 | 0.64 |
| *Ghd7ghd8Hd1* | 38 | 108-116 | -2.0 | NS |  |
| *Ghd7Ghd8Hd1* | 40 | 113-136 | -9.9 | -7.3 | 0.74 |

Negative value indicates the functional allele of *Ghd7.1* promotes rice heading. A, additive effect; D, dominance effect; |D/A|, the degree of dominance; NS, no significance.

### Transcriptional analysis of *Ehd1* and *Hd3a* in the *Ghd7ghd8Hd1, Ghd7Ghd8Hd1* and *Ghd7Ghd8hd1* backgrounds

The florigen gene *RFT1* exhibits a loss of function in ZS97 (Zhao *et al.*, 2015). Considering that *Ghd7.1* has an alternative function in these three backgrounds under different day-length conditions, the expression of *Ehd1* and another florigen gene *Hd3a* was compared between *ghd7.1* and *Ghd7.1* in these three backgrounds under LD and SD conditions, respectively. The relative expression levels of *Ehd1* and *Hd3a* in *Ghd7.1Ghd7ghd8Hd1* and *Ghd7.1Ghd7Ghd8hd1* genotypes decreased under LD conditions but increased under SD conditions compared with those in *ghd7.1Ghd7ghd8Hd1* and *ghd7.1GM7GM8M1*, respectively (Figs 3a-f, S4a-f). The expression of *Ehd1* and *Hd3a* showed no significant difference between *ghd7.1* and *Ghd7.1* in the *Ghd7Ghd8Hd1* background under LD conditions but increased with the presence of *Ghd7.1* under SD conditions (Figs 3b, e; S4b, e). These results indicated that *Ghd7.1* promoted the expression of *Ehd1* and *Hd3a* in these three backgrounds under SD conditions, resulting in an early heading date. In contrast, *Ghd7.1* delayed rice heading in the *ghd7ghd8Hd1* background by repressing the expression of *Ehd1* and *Hd3a* under both LD and SD conditions (Fig. S5a-f).

**Figure 3.**
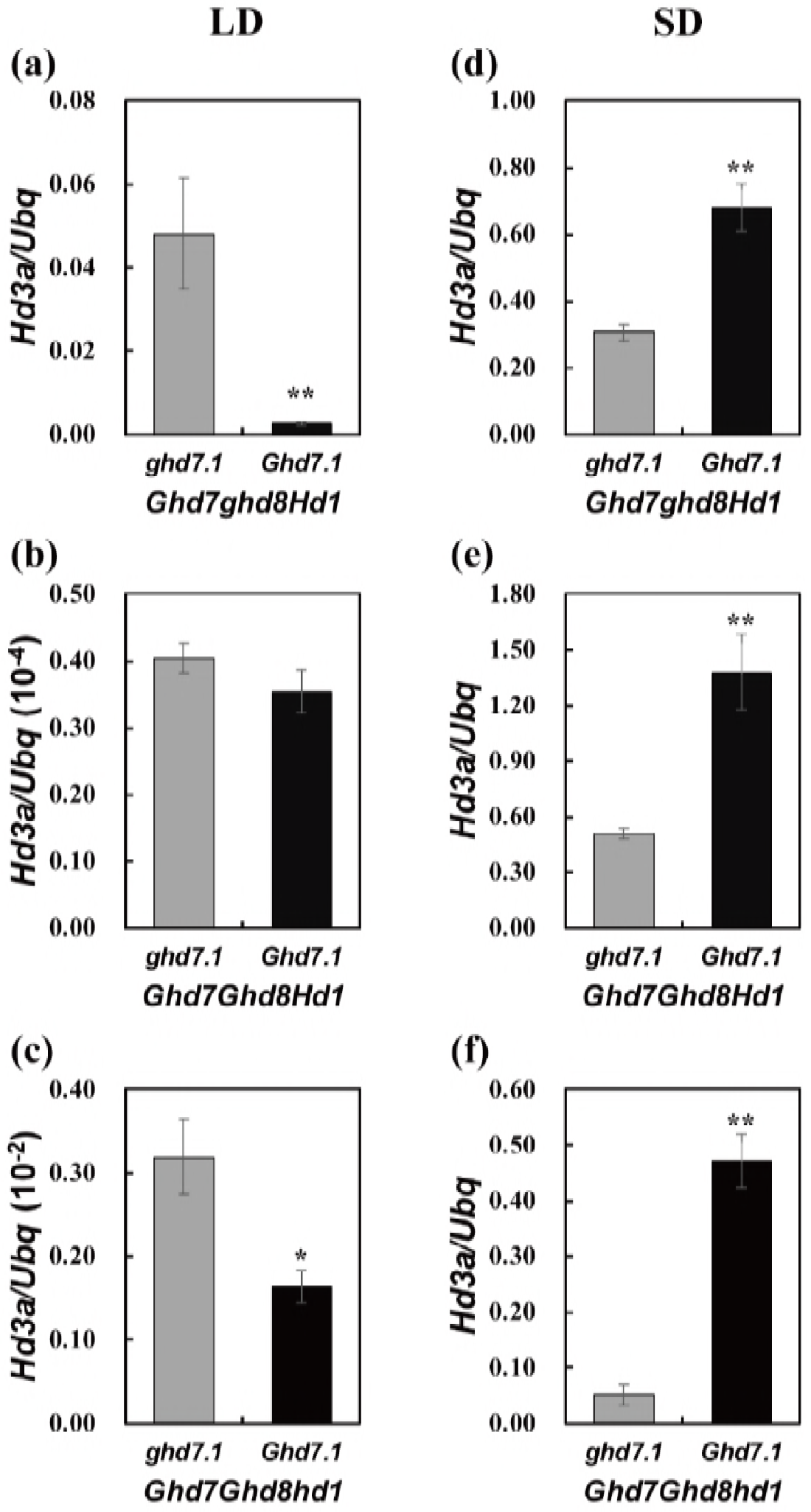
Transcript levels of *Hd3a* in *ghd7.1* and *Ghd7.1* under different day-length conditions. (**a**)-(**c**) Relative expression levels of *Hd3a* between *ghd7.1* and *Ghd7.1* in the Ghd7ghd8Hd1, Ghd7Ghd8Hd1 and *Ghd7Ghd8hd1* backgrounds under LD conditions, respectively; (**d**)-(**f**) Relative expression levels of *Hd3a* between *ghd7.1* and *Ghd7.1* in these three backgrounds under SD conditions, respectively. * and **, *P*<0.05 and *P*<0.01 based on Student’s *t*-test, respectively.

### Correlation between yield and heading date under NLD and NSD conditions

We identified the relationship between heading date and spikelets per panicle (SPP) on the basis of performance of 16 homozygous four-gene combinations. Under NLD conditions, the heading date of these 16 combinations exhibited a continuous distribution ranging from 60 days to 130 days except for the two non-heading combinations *Ghd7Ghd7.1Ghd8Hd1* and *Ghd7ghd7.1Ghd8Hd1*. The earliest heading combination was *ghd7ghd7.1ghd8Hd1*, at 60.8 days, while the latest heading combination in Wuhan was *Ghd7Ghd7.1Ghd8hd1*, at 129.2 days (Fig. 4a). Unexpectedly, SPP of these 14 combinations showed an inverse correlation with heading date. The SPP increased with later heading dates when the heading date was earlier than 90 days, while the SPP decreased with later heading dates when heading date was after 90 days (Fig. 4b). The curve-fitting plots of heading date with SPP under NLD conditions also revealed the inverse correlation with an inflection point at 90.0 days (Fig. 4c). The combination *ghd7Ghd7.1ghd8hd1* had the most SPP, with 199.9±7.3 under NLD conditions, and the second most was *Ghd7ghd7.1ghd8Hd1*, with 184.1±9.8. Under NSD conditions, the heading date of the 16 combinations also showed a continuous distribution with a range from 98 days to 132 days. The combination with the earliest heading date was *ghd7ghd7.1ghd8Hd1*, at 98.7 days, while the combination with the latest heading date was *Ghd7ghd7.1Ghd8hd1*, at 131.8 days (Fig. 4d). The SPP of these 16 combinations increased with the later heading dates, indicating that SPP was positively correlated with heading date under NSD conditions (Fig. 4e, f).

**Figure 4.**
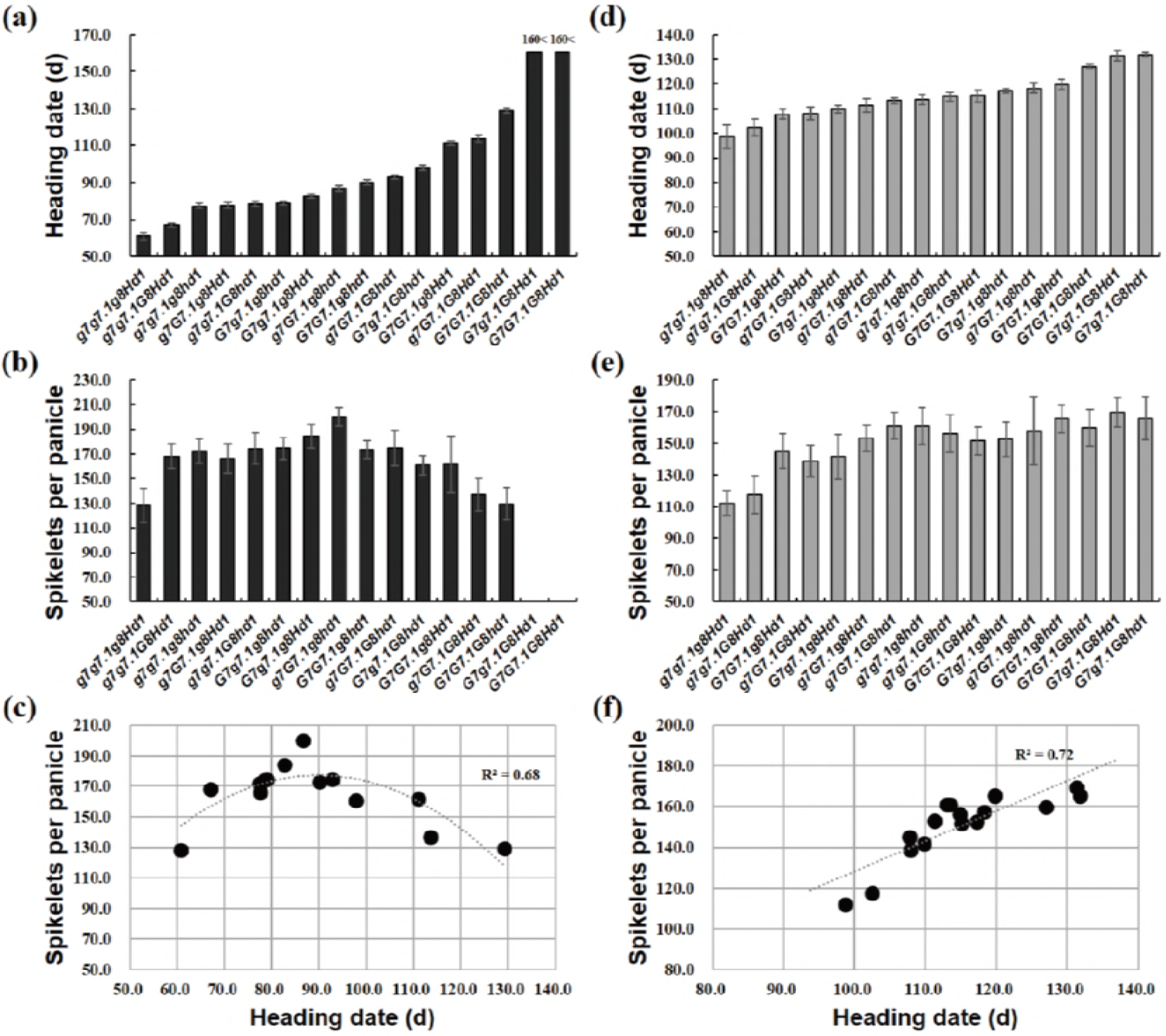
Performances of 16 homozygous combinations of *Ghd7*, *Ghd7.1*, *Ghd8* and *Hd1* on heading date, spikelets per plant under NLD and NSD conditions. Heading date (**a**, **d**), spikelets per plant (**b**, **e**) and Curve-fitting plots of heading date with spikelets per plant (**c**, **f**) under NLD and NSD conditions, respectively. The combinations in (**a**), (**b**), (**d**) and (**e**) are ordered by the increasing heading date. Curves fitting the trait change in (**c**) and (**f**) are calculated by the quadratic and liner equation with *R*^2^ values, respectively. “160<”, no heading after 160 days from sowing. 20≤n≤24 for each combination under NLD conditions and 10≤n≤16 for each combination under NSD conditions.

## Discussion

### *Ghd7*, *Ghd7.1*, *Ghd8* and *Hd1* jointly contributed to a large variation in heading date

*Ghd7*, *Ghd7.1*, *Ghd8* and *Hd1* are all photoperiod sensitive genes that respond to day-length changes and play important roles in rice adaptation to high latitude regions (Yano *et al.*, 2000; Xue *et al.*, 2008; Yan *et al.*, 2011; Liu *et al.*, 2013; Yan *et al.*, 2013). Their combinations also largely determine the adaptation and yield potential of rice cultivars. Loss-of-function allele combination (NNN) and pre-existing strong allele combination (SSF) of *Ghd7*, *Ghd8* and *Hd1* allow rice cultivars to adapt to temperate and tropical regions, respectively (Zhang *et al.*, 2015a). Loss-of-function alleles of *Ghd7*, *Ghd7.1* and *Hd1* contributed to early rice heading dates in the northern regions of northeast China, while functional alleles delayed heading in the southern regions of northeast China, indicating that divergent alleles of these three genes largely determined rice adaptation in northeast China (Ye *et al.*, 2018). In this study, the combinations of *Ghd7*, *Ghd7.1*, *Ghd8* and *Hd1* in ZS97 background exhibited stronger photoperiod sensitivity under NLD conditions than under NSD conditions. Significant digenic, trigenic or even tetragenic interactions of these four genes were detected under both NLD and NSD conditions (Tables 1, 2), but the significance detected under NLD conditions was much greater than that detected under NSD conditions, where the effects of *Ghd7*, *Ghd7.1* and *Ghd8* were decreased. The OsHAPL1-DTH8/Ghd8-Hd1 complex acts as a transcriptional regulator of heading date by interacting with the HAP complex and GTFs (Zhu *et al.*, 2017). *Ghd8/DTH8* encodes a HAP3 subunit, which forms a multicomplex with HAP2 and HAP5 (Thirumurugan *et al.*, 2008). *Ghd7*, *Ghd7.1* and *Hd1* encode transcription factors containing CCT domains, which are similar to HAP2 and responsible for DNA binding and protein-protein interaction (Wenkel *et al.*, 2006; Thirumurugan *et al.*, 2008). Thus, interactions among these genes probably indicate physical interaction among their encoding proteins or between proteins (transcriptional factors) and DNA (gene promoters). In addition, only strong functional and nonfunctional alleles were taken into consideration in this study. The heading date of these 16 four-gene combinations showed a continuous distribution with a range of 60-130 days and no heading under NLD conditions in Wuhan and a range of 98-132 days under NSD conditions in Hainan (Figs S2a, S3a). In nature, there are more diverse alleles for each gene (Koo *et al.*, 2013; Yan *et al.*, 2013; Zhang *et al.*, 2015a). It is expected that different gene combinations will have similar heading dates due to the comprehensive effect of single gene and interaction effects. A better understanding of these four major flowering genes will aid in breeding design for developing cultivars for local rice production. It is noticed that these findings are derived from typical Xian (indica) cultivar, ZS97. It is not clear whether similar results would be obtained in Geng (japonica), which is worth testing in the future.

### Seeking specific four-gene combinations with maximal yield potential for ecological areas

Grain yield is positively correlated with heading date, especially in low latitude areas where the temperature is warm year-round (Gao *et al.*, 2014; Li *et al.*, 2018). In this study, due to continuously high temperature stress during the rice flowering stage in Wuhan, the seed setting rates were significantly decreased; therefore, we analyzed the relationship between heading date and SPP instead of that between heading date and grain yield. The SPP is consistently and positively correlated with heading date under NSD conditions. Nevertheless, The SPP exhibited an inverse correlation with heading date under NLD conditions. The SPP increased with increasing days from sowing to heading when the heading date was earlier than 90 days, while it decreased with increasing days when the heading date was later than 90 days. Based on this finding, optimized combinations can be suggested for local regions to maximize rice production. For example, varieties with the *Ghd7ghd7.1Ghd8Hd1* and *Ghd7ghd7.1Ghd8hd1* combinations will produce more grains in low latitude regions (tropical regions) with short-day and warm conditions such as Hainan. In subtropical regions such as Wuhan, the *ghd7Ghd7.1ghd8hd1* and *Ghd7ghd7.1ghd8Hd1* combinations will have the highest yield potential, at least in the ZS97 background. In this study, the set of materials was grown at only two locations. If they were tested in multiple diverse ecological areas, the favorable gene combinations could be defined for each area.

### Possible mechanism of alternative functions of *Ghd7.1* under NSD conditions

*Ghd7* suppressed the expression of *Hd3a* under LD conditions, whereas it did not affect *Hd3a* expression under SD conditions, resulting in a late heading date in LD but no difference in SD (Xue *et al.*, 2008). Previous studies showed that *Ghd7.1* inhibited heading date under LD conditions but seemed to have no effect under SD conditions (Koo *et al.*, 2013; Liu *et al.*, 2013; Yan *et al.*, 2013; Gao *et al.*, 2014). However, in this study, *Ghd7.1* delayed rice heading in the *ghd7ghd8hd1*, *ghd7ghd8Hd1* and *ghd7Ghd8Hd1* backgrounds but significantly promoted heading in the *Ghd7Ghd8hd1*, *Ghd7Ghd8Hd1* and *Ghd7ghd8Hd1* backgrounds under NSD conditions (Table 4), which clearly demonstrated that *Ghd7.1* had alternative functions under NSD conditions. This differed from the results for *Hd1*, which had alternative functions under NLD conditions (Zhang *et al.*, 2017). *Ghd7.1* suppressed heading date by inhibiting the expression of its downstream genes *Ehd1* and *Hd3a* under LD conditions (Koo *et al.* 2013; Gao *et al.* 2014). *Ghd7.1* acted as an activator of rice heading by promoting *Ehd1* and *Hd3a* expression in the *Ghd7Ghd8hd1*, *Ghd7Ghd8Hd1* and *Ghd7ghd8Hd1* backgrounds under SD conditions (Figs 3, S4). *Ghd7* likely plays an essential role in the function inversion of *Ghd7.1.* However, Ghd7 and Ghd7.1 are both transcriptional suppressors (Weng *et al.*, 2014; Liu *et al.*, 2018). The effect of *Ghd7* on heading date was the largest in the four-gene segregating population under NSD conditions (Table 2). Furthermore, the *Ghd7* by *Ghd7.1* interaction was the strongest digenic interaction in this segregating population under NSD conditions (Tables 2, S3). Consequently, the enhanced genetic interaction between *Ghd7.1* and *Ghd7* under NSD conditions most likely attenuated the interaction of *Ghd7* with other genes, and ultimately upregulated the expression of *Ehd1* and *Hd3a*, resulting in an early heading date.

## Acknowledgments

We would like to thank Mr. Jianbo Wang for his excellent work in the field. This work was supported by the National Key Research and Development Program of China (No. 2016YFD0100301), the National Natural Science Foundation of China (No.31701391, 31701054), and the Natural Science Foundation of Hubei Province (No.2015CFA006).

## Author contributions

Y.X., H.L. and B.Z. planned and designed the research. B.Z. performed most of experiments and conducted fieldwork. B.Z., H.L. and Z.Z. prepared the materials. F.Q. and Q.L. performed QTLs genotyping. B.Z. and Z.H. analyzed data. B.Z. and Y.Z. wrote and revised the manuscript.

## Supplemental material

**Figure S1.**
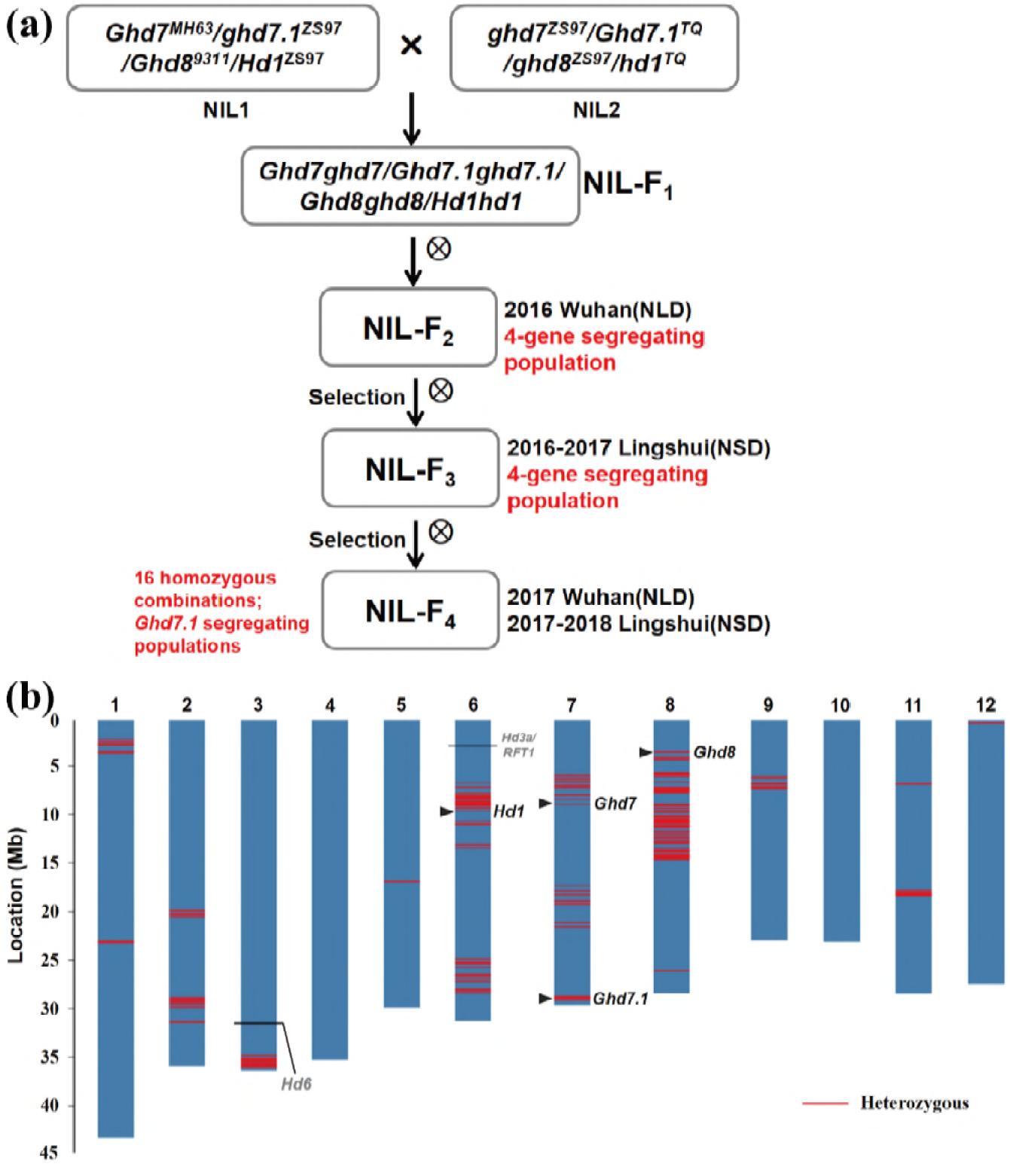
Development and genome composition of the rice populations. (**a**) Development of the near-isogenic population segregating for 4 heading date genes by crossing NIL1 with NIL2. NIL-1 was introgressed with functional *Ghd7*^*MH63*^ and *Ghd8*^*9311*^ in ZS97. NIL-2 was introgressed with functional *Ghd7.1*^*TQ*^ and nonfunctional *hd1*^*TQ*^ in ZS97. Thus, the NIL-F_1_ plants contained heterozygous *Ghd7*, *Ghd7.1*, *Ghd8* and *Hd1*. (**b**) Visualization of the genome composition of the NIL-F_1_ plant obtained by using the Rice6K SNP array. The physical positions of *Ghd7*, *Ghd7.1*, *Ghd8* and *Hd1* are marked by black triangles. The homozygous *Hd6* on the third chromosome and *Hd3a/RFT1* on the sixth chromosome are marked by black lines. Red lines indicate heterozygous chromosomal regions based on SNP markers.

**Figure S2.**
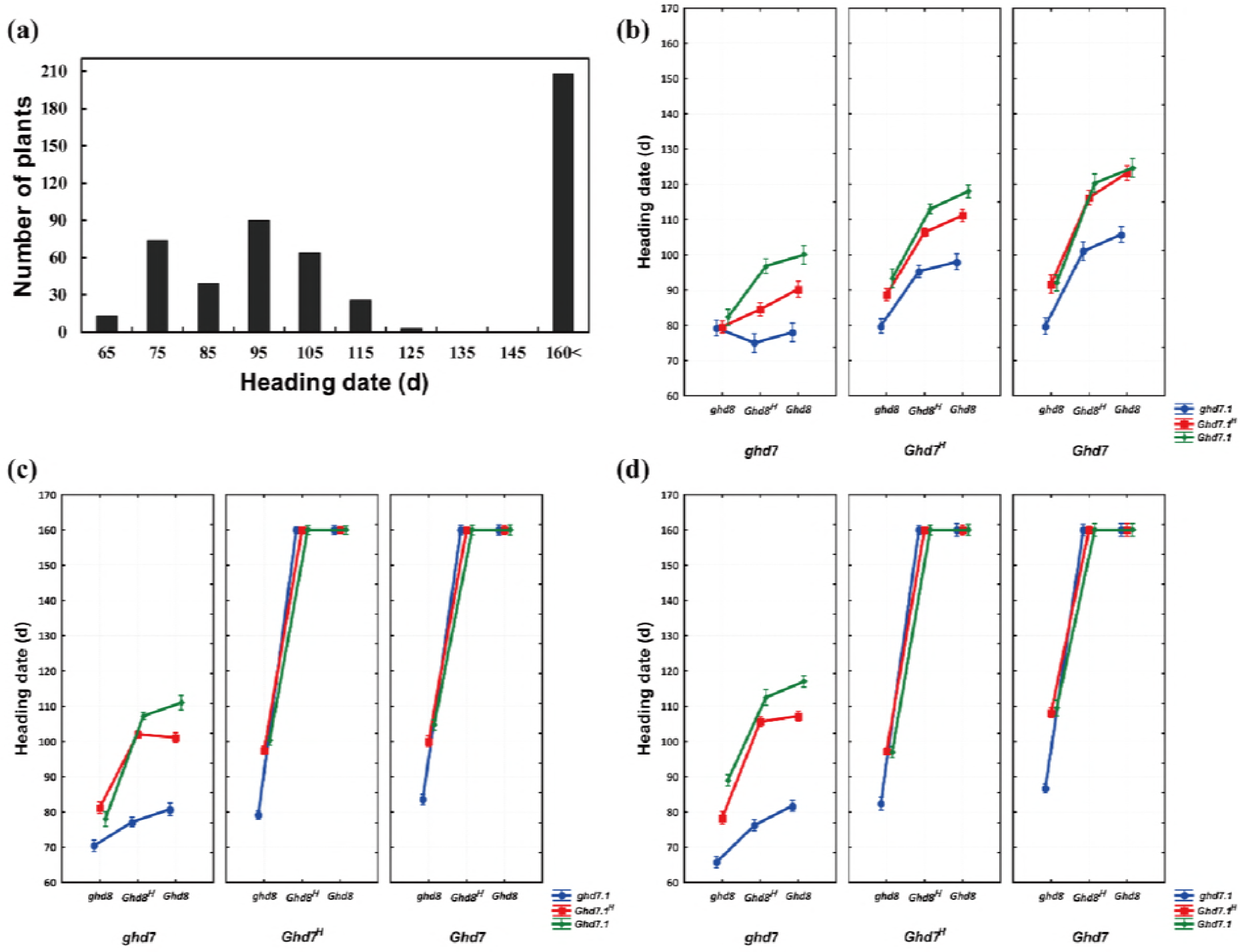
Genetic interaction analysis of heading date genes in the NIL-F_2_ population under NLD conditions. (**a**) The distribution of heading date. Three-way ANOVA of *Ghd7*, *Ghd7.1* and *Ghd8* in the *Hd1* (**b**), *Hd1*^*H*^ (**c**) and *hd1* (**d**) backgrounds. *Ghd7*^*H*^, *Ghd8*^*H*^ and *Ghd7.1*^*H*^ indicate the heterozygous alleles of *Ghd7, Ghd8* and *Ghd7.1*, respectively. Data are represented by LS means. Vertical bars denote 0.95 confidence intervals.

**Figure S3.**
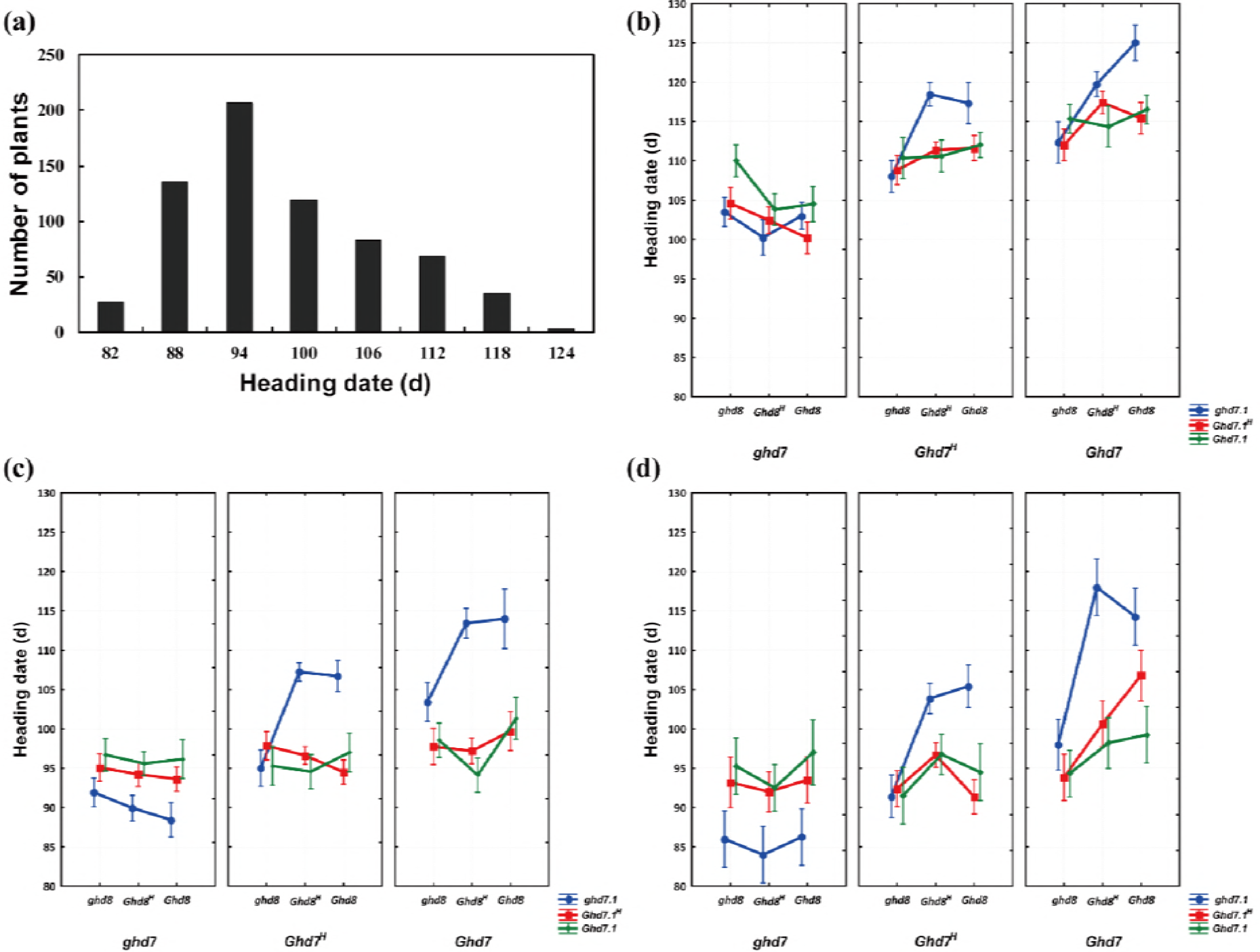
Genetic interaction analysis of heading date genes in the NIL-F_3_ population under NSD conditions. (**a**) The distribution of heading date. Three-way ANOVA of *Ghd7*, *Ghd7.1* and *Ghd8* in the *Hd1* (**b**), *Hd1*^*H*^ (**c**) and *hd1* (**d**) backgrounds. *Ghd7*^*H*^, *Ghd8*^*H*^ and *Ghd7.1*^*H*^ indicate the heterozygous alleles of *Ghd7*, *Ghd8* and *Ghd7.1*, respectively. Data are represented by LS means. Vertical bars denote 0.95 confidence intervals.

**Figure S4.**
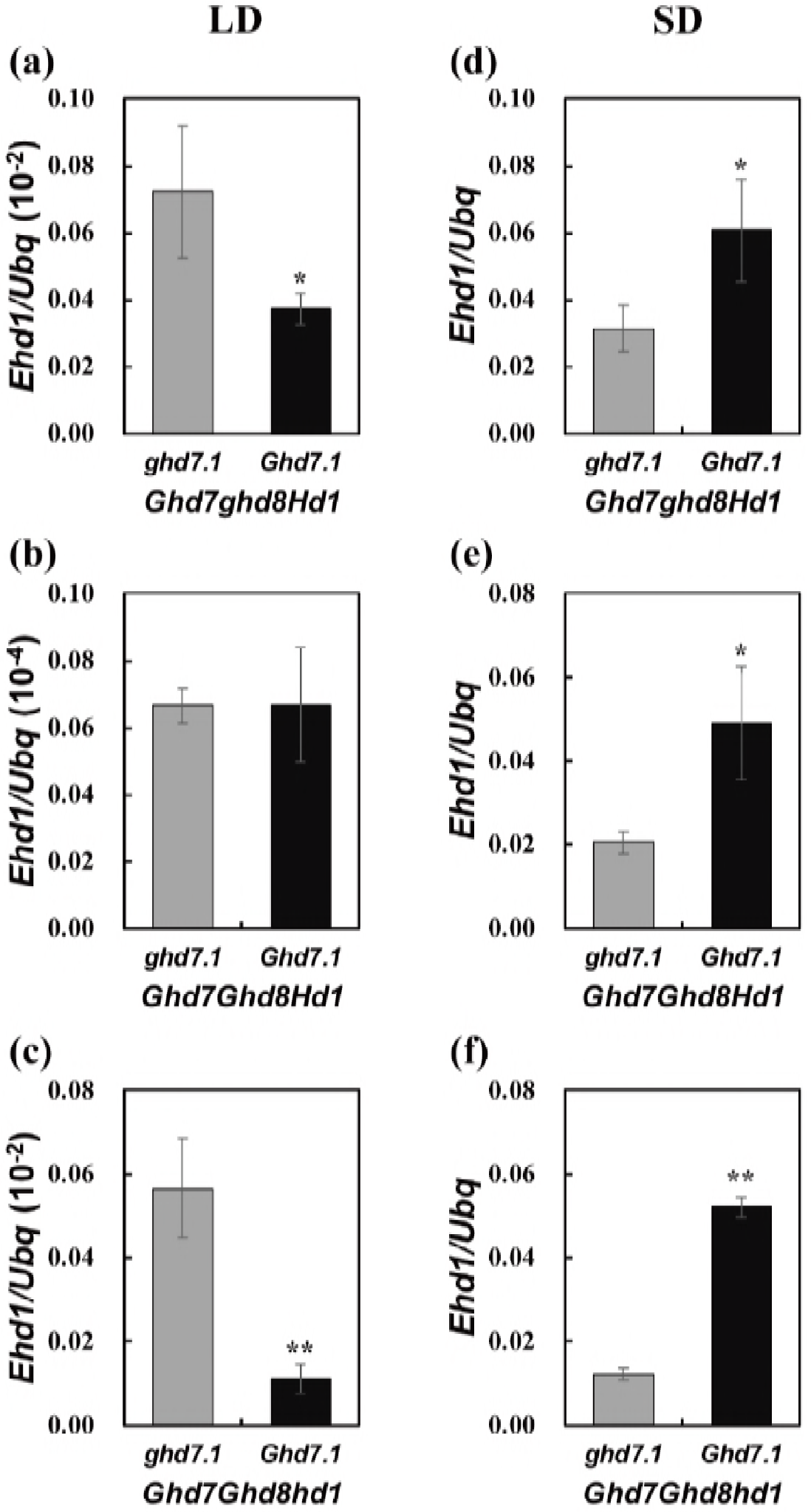
Transcript levels of *Ehd1* in *ghd7.1* and *Ghd7.1* under different day length conditions. (**a**)-(**c**) Relative expression levels of *Ehd1* between *ghd7.1* and *Ghd7.1* in the *Ghd7ghd8Hd1*, *Ghd7Ghd8Hd1* and *Ghd7Ghd8hd1* backgrounds under LD conditions, respectively; (**d**)-(**f**) Relative expression levels of *Ehd1* between *ghd7.1* and *Ghd7.1* in these three backgrounds under SD conditions, respectively. * and **, *P*<0.05 and *P*<0.01 based on Student’s *t*-test, respectively.

**Figure S5.**
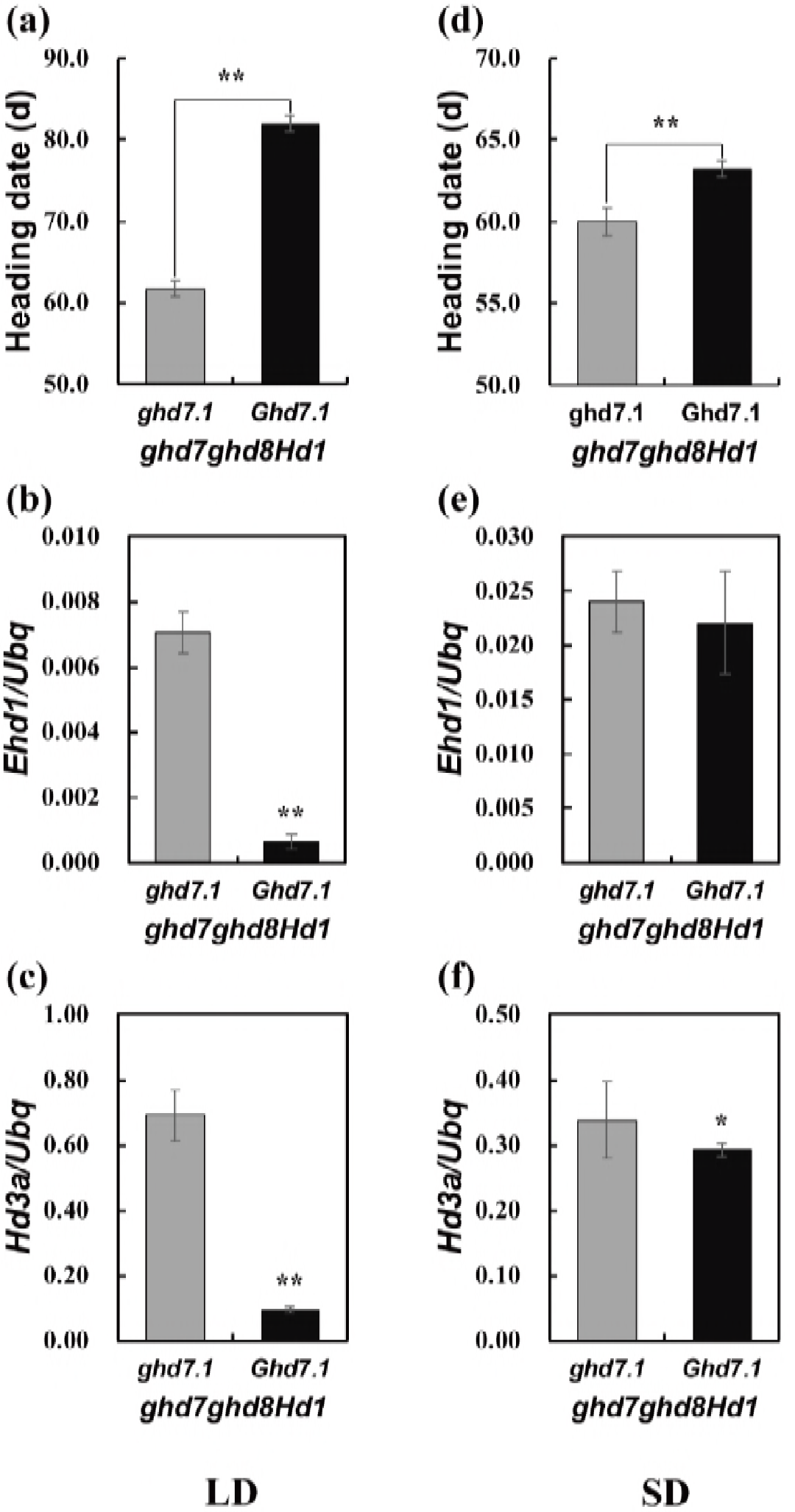
*Ghd7.1* delays the heading date in the *ghd7ghd8Hd1* background under both LD and SD conditions. Heading date (**a**), *Ehd1* expression (**b**) and *Hd3a* expression (**c**) in *ghd7.1* and *Ghd7.1* in the *ghd7ghd8Hd1* background under LD conditions; Heading date (**d**), *Ehd1* expression (**e**) and *Hd3a* expression (**f**) in *ghd7.1* and *Ghd7.1* in the *ghd7ghd8Hd1* background under SD conditions. * and **, *P*<0.05 and *P*<0.01 based on Student’s *t*-test, respectively.

**Table S1.** Characteristics of four heading date genes and linked markers.

| Gene | Varieties donated alleles |  | Markers* | Reference |
| --- | --- | --- | --- | --- |
|  | Non-functional | Functional |  |  |
| *Ghd7* | ZS97 (Completely absent) | MH63 | MRG4436 | Xue *et al.* 2008 |
| *Ghd7.1* | ZS97 (Frame shift, 8bp deletion) | Teqing | InDel37 | Yan *et al.* 2013 |
| *Ghd8* | ZS97 (1116-bp deletion in 3' region) | 9311 | Z9M | Yan *et al.* 2011 |
| *Hd1* | Teqing (Frame shift, 4bp deletion) | ZS97 | S56 | Zhang *et al.* 2017 |

\* MRG4436 is a marker tightly linked to *Ghd7*, and the other three markers are functional markers derived from the corresponding genes.

**Table S2.**
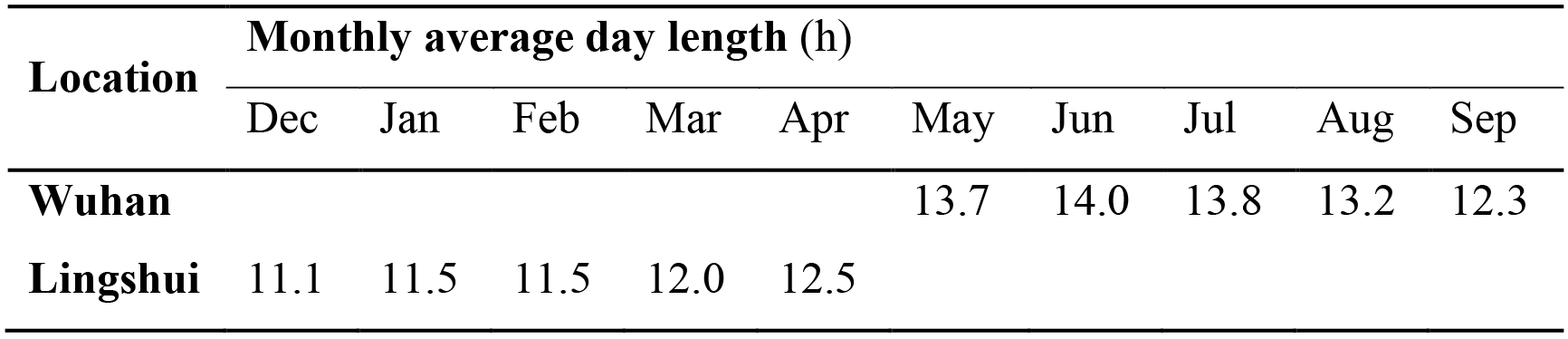
The monthly average day length of growing seasons at Wuhan and Lingshui.

| Location | Monthly average day length (h) |  |  |  |  |  |  |  |  |  |
| --- | --- | --- | --- | --- | --- | --- | --- | --- | --- | --- |
|  | Dec | Jan | Feb | Mar | Apr | May | Jun | Jul | Aug | Sep |
| **Wuhan** |  |  |  |  |  | 13.7 | 14.0 | 13.8 | 13.2 | 12.3 |
| **Lingshui** | 11.1 | 11.5 | 11.5 | 12.0 | 12.5 |  |  |  |  |  |

**Table S3.** The paired digenic interactions between *Ghd7*, *Ghd7.1*, *Ghd8* and *Hd1* in the four-gene segregating population under NLD and NSD conditions.

| Effect | DF | NLD (n=509) |  | NSD (n=679) |  |
| --- | --- | --- | --- | --- | --- |
|  |  | F | P | F | P |
| *Ghd7* by *Ghd7.1* | 4 | 150 | <1.0E-10 | 94.8 | <1.0E-10 |
| *Ghd7* by *Ghd8* | 4 | 1436 | <1.0E-10 | 36.5 | <1.0E-10 |
| *Ghd7* by *Hd1* | 4 | 1078 | <1.0E-10 | 10.6 | 2.5E-08 |
| *Ghd7.1* by *Ghd8* | 4 | 9 | 4.8E-07 | 27.9 | <1.0E-10 |
| *Ghd7.1* by *Hd1* | 4 | 16 | <1.0E-10 | 4.6 | 1.1E-03 |
| *Ghd8* by *Hd1* | 4 | 1296 | <1.0E-10 | 8.1 | 2.4E-06 |

DF, degree of freedom.

**Table S4.**
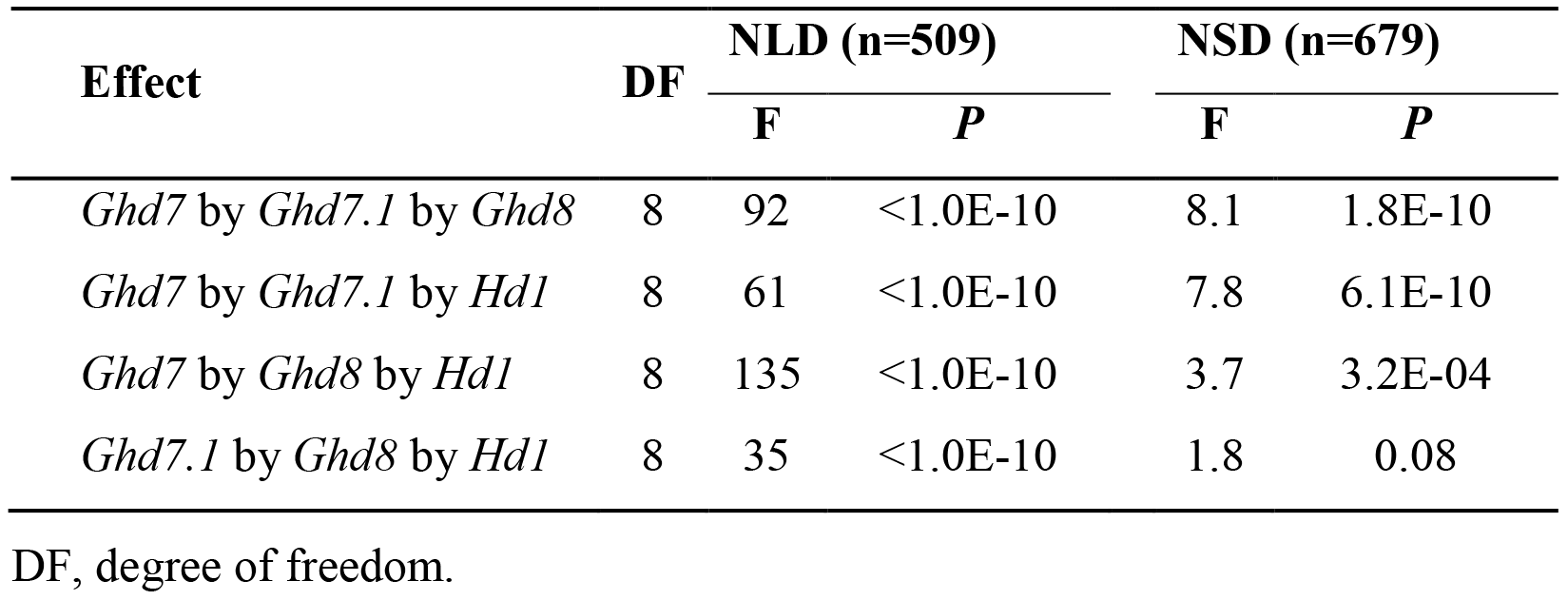
The trigenic interactions among *Ghd7*, *Ghd7.1*, *Ghd8* and *Hd1* in the four-gene segregating population under NLD and NSD conditions.

**Table S5.**
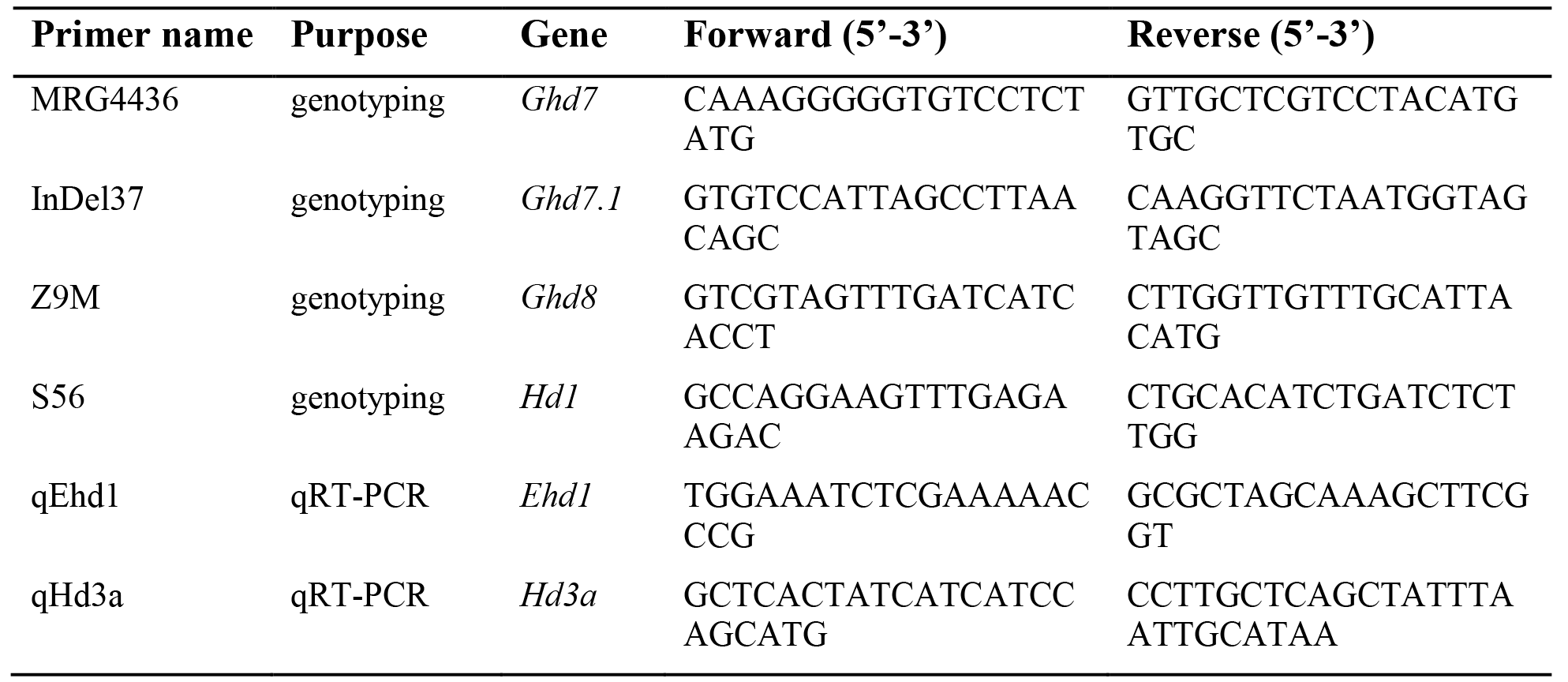
Primers used in this study.

